# Sleep-like slow waves in wakefulness index post-stroke behavioural alterations

**DOI:** 10.1101/2025.11.24.689850

**Authors:** Daniel J. Pearce, Mana Biabani, Méadhbh B. Brosnan, Trevor T.-J. Chong, Nele Demeyere, Ger M. Loughnane, Jason B. Mattingley, Margaret J. Moore, Peter W. New, Dragan Rangelov, Redmond G. O’Connell, Renerus J. Stolwyk, Sam S. Webb, Thomas Andrillon, Mark A. Bellgrove

## Abstract

Increases in high-amplitude, low-frequency EEG oscillations during non-rapid eye movement (NREM) sleep serve as markers of neuronal silencing, which disrupts functional connectivity and is associated with a loss of awareness and responsiveness. Cortical slowing likewise occurs during wakefulness, upon sleep deprivation, and accompanies performance deterioration. Recent work has revealed similar oscillations (termed ‘sleep-like slow waves’) following stroke in brain regions close or connected to a brain lesion, offering a plausible but untested mechanism for stroke-related behavioural impairment. Here, we simultaneously recorded EEG and behavioural performance while 30 unilateral stroke patients (12 left hemisphere, 18 right hemisphere) and 27 age-matched healthy adults completed a computerised perceptual decision-making task. Sleep-like slow waves occurred more frequently in stroke patients relative to healthy controls. Furthermore, there was a significant correlation between slow wave occurrence and the accuracy and speed of perceptual reports in stroke patients, suggesting that variations in the occurrence of slow waves may contribute to cognitive heterogeneity post-stroke. Evidence that sleep-like slow waves predict deficits in perceptual decision-making following stroke indicates that stroke-induced lesions can destabilize cortical networks extending beyond the damaged regions, thereby contributing to broader cognitive impairments.

## Introduction

Cognitive impairment after stroke is highly prevalent, with estimates varying depending on time post-stroke, the measures used and the cohort studied (e.g. (Jokinen et al., 2015; Pendlebury & Rothwell, 2019; Sexton et al., 2019)). When assessed with a highly sensitive tool, the majority of acute stroke patients show a cognitive deficit, which is unsurprising given the acute brain injury (Webb et al., 2022). Whilst many patients recover fully, long-term cognitive impairments persist in 30-50% of stroke survivors (Barker-Collo et al., 2010; Kusec et al., 2023) including poorer quality of life and everyday functioning (Stolwyk et al., 2021, 2024). In particular, attentional deficits including a reduced ability to respond to contralesional stimuli may remain detectable long after the stroke (e.g., (Losier & Klein, 2001; Pearce et al., 2023)). Stroke patients also experience excessive daytime sleepiness and mental fatigue (Ding et al., 2016; Johansson & Rönnbäck, 2012). However, long-term outcomes are markedly heterogeneous, and we lack reliable objective predictors of stroke recovery (Price et al., 2017). Fluctuations in cortical arousal, i.e. the level of excitability of the cortex, could be a major source of post-stroke outcome heterogeneity, affecting behaviour, cognition, and emotion (Calderon et al., 2016; Eason et al., 1969; Petersen et al., 2017; Pfaff et al., 2008; Pfaff & Banavar, 2007; Pfaff & Kieffer, 2008). Indeed, the severity of post-stroke perceptual difficulties is known to be exacerbated by low arousal (Lazar et al., 2002; Robertson et al., 1998), potentially arising from disruptions of the regulation of cortical excitability (Chaudhuri & Behan, 2004; Chen et al., 2023).

States of low cortical arousal, e.g. induced by sleep pressure or anaesthetics, can be tracked with scalp EEG, and are characterised by a shift of brain dynamics toward slower frequencies (i.e., in the delta or theta ranges, 1-7 Hz) (Murphy et al., 2011; Purdon et al., 2013). This shift has been interpreted to reflect the intrusion of locally generated sleep-like dynamics (or ‘slow waves’) (Colombo et al., 2025), similar to those seen in non-rapid eye movement (NREM) sleep (Sarasso et al., 2020; Vyazovskiy & Harris, 2013) but occurring within a global states of wakefulness, a phenomenon termed local sleep (Andrillon et al., 2019; Andrillon & Oudiette, 2023; Krueger et al., 2019). Stroke has been well-documented to also result in increased power in slower frequencies, particularly in regions near to the lesion (Murri et al., 1998; Nuwer, 1996; Yokoyama et al., 1996). As for sleep deprivation, recent work has attributed this cortical slowing to the intrusion of transient and localised high amplitude, low frequency (1-7 Hz) slow waves (SWs). This increase is predominant in brain regions close or connected with the lesioned area (Massimini et al., 2024), possibly due to changes in excitatory-inhibitory balance or effective connectivity. Importantly, studies conducted in animals (Rector et al., 2005, 2009; Vyazovskiy et al., 2011) and humans (Avvenuti et al., 2021; Bernardi et al., 2015; Hung et al., 2013) showing the occurrence of SWs in healthy but sleep-deprived or fatigued individuals established a region and task-specific correlation between SW occurrence and behavioural impairment (e.g., errors, misses, false alarms, and slow responses), suggesting that such SWs could have a direct impact on cognitive processes.

Thus, the measurement of SWs with non-invasive EEG recordings could not only help to map the cortical networks impacted by the stroke (Massimini et al., 2024), but it could also offer mechanistic insights into the source of cognitive heterogeneity post-stroke. Indeed, sleep SWs recorded with scalp or depth EEG represent, at the neuronal level, so-called OFF periods or brief episodes of reduced or absent neural firing (Steriade et al., 1991; Vyazovskiy et al., 2009). These OFF periods disrupt cortical processing and cortico-cortical interactions ultimately leading to the loss of awareness and responsiveness typically associated with sleep (Andrillon & Kouider, 2019; Massimini et al., 2007; Rosanova et al., 2018; Tononi & Massimini, 2008). This relationship between SWs and reduced neuronal firing seems to extend to SWs in wakefulness (Marmelshtein et al., 2023; Sheybani et al., 2023; Vyazovskiy et al., 2011), offering a simple mechanistic account of how the occurrence of sleep-like SWs in wakefulness could impact behavioural performance (Andrillon et al., 2019). SWs could also play a protective role following stroke, in line with the restorative function of sleep SWs (Krueger et al., 2019; Léger et al., 2018; Vyazovskiy & Harris, 2013) as increased low frequency activity in the acute stages post-stroke is predictive of better outcomes (Finnigan et al., 2004; Sarasso et al., 2020).

In this study, we investigated the relationship between the presence of sleep-like SWs following stroke and performance on a perceptual decision-making task that examines the dynamic integration of perceptual information to make an appropriate decision. We hypothesised that SWs would occur more frequently in stroke patients compared with age-matched controls, with a higher density in the lesioned hemisphere, and that SW density would correlate negatively with behavioural performance. Further, we expected a relatively higher rate of sleep-like SWs to correlate with poorer self-reported everyday functioning and greater reports of post-stroke fatigue.

## Materials & Methods

### Participants

Behavioural and EEG data were collected from 27 healthy adults aged between 57 and 90 years (*M_age_* = 73.44 years, *SD_age_* = 7.06; *n* = 13 male, *n* = 14 female) and 30 stroke patients aged between 53 and 84 years (*n* = 12 left hemisphere stroke, *n* = 18 right hemisphere stroke; *M_age_* = 70.40 years, *SD_age_* = 7.99; *n* = 16 male, *n* = 14 female) while they completed a bilateral variant of the random dot motion task (Fig. 1) (Newsome et al., 1989; Britten et al., 1992; Loughnane et al., 2016; Pearce et al., 2025). Data collection took place in two centres: Monash (Australia) (N=27 healthy controls; N=11 stroke patients) and Oxford (United Kingdom) University (N=19 stroke patients). All participants were right-handed, reported normal or corrected-to-normal visual acuity, and were not taking arousal-modulating medications. Healthy adults reported no prior history of neurological or psychiatric disease and demonstrated no frank cognitive impairment, as operationalised by a score of over 23/30 on the Montreal Cognitive Assessment (MoCA) (Nasreddine et al., 2005) after adjusting for education as per (Carson et al., 2018). The stroke patients had a medically confirmed diagnosis of first clinically significant, unilateral stroke without any other neurological or psychiatric diagnoses and did not have inferior visual field defects (stimuli were presented to the inferior quadrants, as ensured by eye tracking-enforced central fixation). Further details and lesion data of stroke patients are reported in (Pearce et al., 2023).

**Figure 1.**
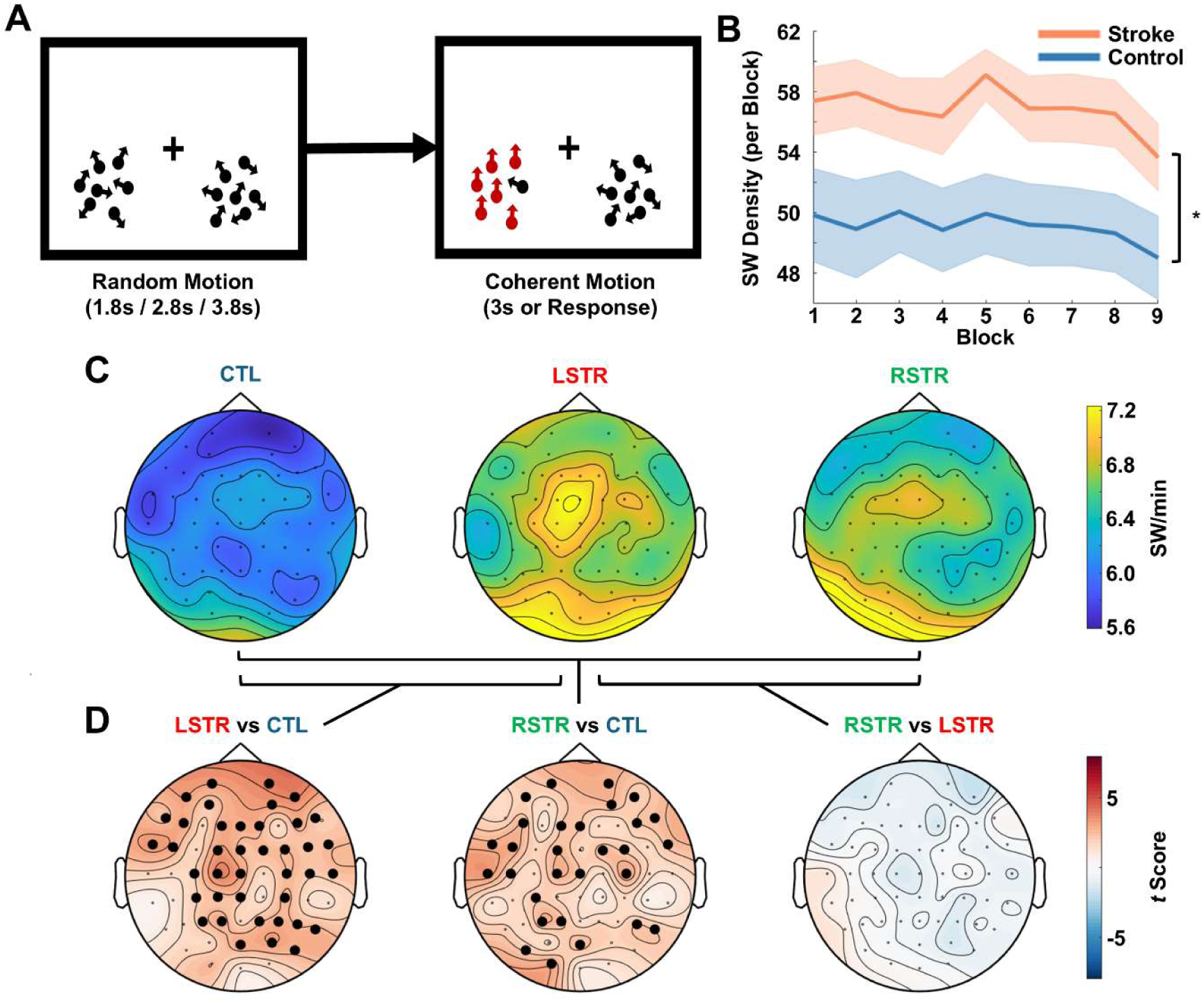
Slow Wave (SW) Density Increases Following Stroke. *Note.* (A) Bilateral Random Dot Motion (BRDM) task. Following central fixation, dots in circular patches either side of the vertical midline began to move randomly. After 1.8, 2.8, or 3.8s, 90% of the dots in one of the two patches moved coherently, either up or down, to which participants responded with a single mouse click. (B) Slow wave (SW) density per block across all blocks, depicted separately for controls and patients. \**p*<.05. (C) Topographical maps of mean SW density per minute, presented separately for controls (CTL; left), left hemisphere (LSTR; centre), and right hemisphere stroke patients (RSTR; right). SW/min = SW per minute. (D) Topographical maps of *t*-values denoting statistical differences between left-hemisphere stroke and controls (left), right-hemisphere stroke and controls (centre), and right- and left-hemisphere stroke (right). Electrodes that form part of significant clusters (*p_cluster_*<.05) are denoted by bolded dots.

### Procedure & Materials

#### Bilateral Random Dot Motion Task

Participants were seated alone at a viewing distance of 57cm in a darkened and sound-attenuated room, with their head supported by a chin rest. The task was run on a computer running MATLAB (MathWorks) and the Psychophysics Toolbox extensions (Brainard, 1997; Cornelissen et al., 2002; Pelli, 1997) for Windows XP. Instructions and visual stimuli were displayed on either a 21-inch CRT monitor, or a 23- or 27-inch LED monitor, depending on the recruitment site. The task was however presented at a consistent screen resolution of 1024 x 768 with an 85Hz refresh rate. Task instructions were provided verbally, and participants completed practice trials until their understanding of the task requirements was deemed sufficient. Note that perfect performance was not required to progress to the main task.

After the training and during the task, participants were required to fixate on the centre of the screen and encouraged to minimise blinking and moving during each trial, with compliance monitored with an EyeLink 1000 (SR Research Ltd) eye tracking camera. While maintaining central fixation they simultaneously monitored two peripheral circular patches presented in the left and right inferior quadrants of the screen (Fig. 1). Each patch encompassed 150 9x9 pixel dots, which moved randomly at a speed of five degrees per second. At set periods of time, 90% of the dots in one of the two patches began to move uniformly in an upward or downward direction. Participants were required to detect the onset of coherent motion, irrespective of direction, and respond via a speeded button press with their right index finger (*n* = 2 used their left index finger due to right-sided hemiplegia) as quickly as possible without making errors. In the event of a blink or a gaze deviation more than four degrees left or right of centre, the dots temporarily ceased moving until fixation was restored and the trial restarted.

The task proceeded for between 7-15 blocks (dependent on participant capacity) of 24 trials each block, with blocks interspersed by breaks of 60-120 seconds. Each trial began at central fixation and encompassed a period of random motion of either 1.8, 2.8 or 3.8 seconds, followed by coherent upward or downward motion in one of the two patches. This gave rise to 12 possible trial types (a combination of three temporal lengths of random motion, two target locations and two coherent motion directions) which were presented in a pseudorandom order. Each trial type was presented twice per block. Motion ceased following a response or after 3 seconds in the event of a non-response. We analysed data from blocks one to nine only, as fewer participants performed more than nine blocks.

#### Behavioural Data

We defined each trial as the time between two instances of central ocular re-fixation as monitored by an eye tracking camera and derived three types of trial: fixation break (i.e., those that were restarted as the participant did not maintain central fixation), miss (i.e., trials with either no response, or one that fell outside the 150-3000ms post-coherent motion onset window), and hit (i.e., trials with a response between 150-3000ms post-coherent motion onset). We excluded fixation break trials from further analysis and then calculated subject-level hit rate by dividing the number of remaining trials that were responded to per block by the total number of remaining trials. For those trials with a response, we calculated the time from coherent motion onset to response (response time, RT) in milliseconds.

### Electroencephalography *(EEG)*

Continuous high-density EEG was acquired from 64 scalp electrodes throughout the task using either a BrainAmp DC system (Brain Products) digitised at 500Hz (Monash) or a Neuroscan SynAmpsRT system digitised at 1000Hz (Oxford), depending on testing site. We processed the data using custom scripts (available at https://github.com/andrillon/LS_Stroke) in combination with FieldTrip (Oostenveld et al., 2011) routines implemented in MATLAB (MathWorks).

#### Pre-Processing

First, we accounted for cross-site differences in data collection by identifying a common subset of 59 electrodes, and down-sampling data collected at Oxford from 1,000 Hz to 500 Hz. All data were detrended and subjected to 50, 100, and 150 Hz notch filters to eliminate line noise, followed by a 0.1 Hz Hamming-windowed FIR high-pass filter. We then applied a 120 Hz low-pass filter, and re-referenced the data to the average reference. Finally, using FastICA we applied an automated Independent Component Analysis (ICA) procedure to remove artifacts related to ocular and muscular movement (Hyvärinen & Oja, 2000; Rogasch et al., 2014). Components associated with ocular and muscular activity were visually selected.

#### Slow Wave (SW) Detection

We extracted sleep-like SWs from continuous EEG data using previously established procedures (Andrillon et al., 2021; Pinggal et al., 2022) adapted from those used to identify SWs during sleep (Riedner et al., 2007). The EEG data were segmented over each block of the task, re-referenced to the average of the TP7 and TP8 channels (closest electrodes to the left and right mastoid bones in this montage), de-trended by removing the average amplitude over each block and resampled to 128 Hz, following which we applied a 1-10Hz Chebyshev type-II bandpass filter. We then identified all zero-crossings in the time series data and defined individual waves as the activity between two negative zero-crossings (i.e., the beginning and end of the wave) and extracted the duration and peak-to-peak (positive – negative peaks) amplitude of each individual wave. The wave frequency was defined as the inverse of the duration. Consistent with previous work focusing on the low frequency range (encompassing the δ [1-4 Hz] and θ [4-7 Hz] bands (Andrillon et al., 2021; Bernardi et al., 2015; Hung et al., 2013; Pinggal et al., 2022), we discarded waves with a frequency over 7 Hz. Additionally, waves that had a positive peak over 75 µV or that occurred within 1 second of a high-amplitude event (with an absolute amplitude of >150 µV) were removed. Finally, we isolated the largest SWs by setting a per-subject, per-block threshold at the 90^th^ percentile of peak-to-peak amplitude separately for each electrode (threshold: *M_control_* = 17.86 µV, *SD_control_* = 8.28, 95% CI_control_ = [17.73,17.99]; *M_LStr_* = 17.53 µV, *SD_LStr_* = 7.15, 95% CI*_LStr_* = [17.36,17.70]; *M_RStr_* = 16.77 µV, *SD_RStr_* = 7.56, 95% CI*_RStr_* = [16.62,16.92]). Thus, we selected the top 10% of SWs with the highest amplitude at each electrode for each subject, controlling for individual subject- and channel-level differences in baseline voltage.

For each of the supra-threshold SWs, we computed the density (number of SW events per minute per electrode per subject), peak-to-peak amplitude (different of amplitude between the most positive and most negative peak of the wave), oscillation frequency (1/wave duration), downward slope (gradient of the negative component of the wave from the onset (zero-crossing) to the negative peak), and upward slope (gradient of the positive component of the wave from the negative peak to the following positive peak) consistent with previous publications (Andrillon et al., 2021; Pinggal et al., 2022).

Additionally, to visualise between-group differences in SWs, we derived an event-related potential of the average SW. To do so, we time-locked the pre-processed EEG data to the onset (first negative zero-crossing) of each SW and aggregated the waveform across a Region of Interest (ROI; outlined below) separately for controls and patients.

### Questionnaires

#### Fatigue Severity Scale (FSS)

We used the FSS (Krupp, 1989) to estimate self-reported fatigue severity in stroke patients only. Participants rated their agreement with nine statements about fatigue (e.g. ‘I am easily fatigued’ and ‘fatigue causes frequent problems for me’) on a seven-point Likert scale from Strongly Disagree to Strongly Agree. Scores therefore range from 9-63, with higher scores indicative of greater fatigue severity. The FSS shows strong psychometric properties in measuring post-stroke fatigue (Nadarajah et al., 2017; Ozyemisci-Taskiran et al., 2019).

#### Patient Competency Rating Scale (PCRS)

The PCRS (Prigatano & Neuropsychological Rehabilitation Program (Presbyterian Hospital, 1986) is a 30-item self-report scale that has been validated for use in estimating everyday psychosocial functioning after a stroke (Barskova & Wilz, 2006). Stroke patients rated the degree of difficulty with which they could perform everyday tasks (e.g., dressing themselves and remembering their daily schedule) on a five-point Likert scale from 1 (“Can’t do”) to 5 (“Can do with ease”). Summed total scores range from 30-150, with higher scores indicative of higher functioning.

### Statistical Analysis

To examine the effect of the variables of interest (within-subject: block effect; between-subject: group effect), we implemented a model comparison approach using linear mixed effect models (LME). We provide below examples of how this strategy was implemented.

For example, we checked that the threshold used to detect SWs did not differ between patients and controls (hereafter referred to as group), since this threshold was adjusted at the subject level (see Results section). To do so, we compared a model with a fixed effect of block and electrode (random effect of subject) with a model with a fixed effect of block, electrode and group. The two models were compared with a Likelihood Ratio Test. This comparison indicated that the model with the fixed effect of group did not lead to a better fit (Likelihood Ratio Test: 0.024; p-value = 0.88). The corresponding model also showed no effect of group (F-value=0.02; p-value=0.88). This analysis indicates that we can compare the supra-threshold SWs across groups.

We also examined the influence of block, electrode location and group on SW density by relying on a model comparison approach. We fitted the following five models:

- Model 1: SW Density ∼ 1 + Electrode + (1|SubID)
- Model 2: SW Density ∼ 1 + Electrode + Block + (1|SubID)
- Model 3: SW Density ∼ 1 + Electrode * Block + (1|SubID)
- Model 4: SW Density ∼ 1 + Electrode * Block + Group + (1|SubID)
- Model 5: SW Density ∼ 1 + Electrode * Block * Group + (1|SubID)

Models of increasing complexity were compared pairwise from the simplest (Model 1) to the most complex (Model 5). Except for Model 5, increasing the complexity of the model significantly improved the fit of the data (Likelihood Ratio Test; all p < 0.05). We report the outputs of Model 4 in the Results section. Of note, for this and subsequent analyses, we corrected for a possible confound of block duration on arousal-related phenomena such as slow-wave activity by excluding blocks longer than three standard deviations above the overall mean (*M*=2.92 minutes, *SD*=0.95; 12 blocks in total across five participants).

To examine the spatial distribution of the effects, we fitted a SW Density ∼ Group + Block + (1|SubID) linear mixed-effects model (LMM) separately at each electrode. In order to account for the multiplication of dependent statistical tests, we used a cluster-permutation approach (Maris & Oostenveld, 2007). We first separated the stroke patients into those with left hemisphere (n=12) and right hemisphere strokes (n=18) to examine whether the distribution of SW density differed depending on the lesioned hemisphere. We then computed the *t*- and *p*-values for each relevant group comparison (control vs left stroke, control vs right stroke, and left vs right stroke), and defined clusters based on neighbouring electrodes that had *p*-values less than the cluster alpha of .05. The *t*-value of each electrode within a given cluster was summed to give the cluster *t*-value, which was then compared to the distribution of cluster *t*-values that resulted when the procedure was repeated with permuted datasets (*N*=500 permutations). Each permuted dataset resulted from randomly shuffling the group identity of each datapoint, with identical shuffling used for each permutation across all electrodes. Electrodes within clusters with a resultant Monte Carlo *p*-value below .05 were deemed statistically significant.

To investigate the properties of SWs following stroke, we defined a Region of Interest (ROI) by repeating the above procedure with a contrast between controls and patients (combining both left-and right-hemisphere stroke groups as no significant differences were found between these two groups in the topographical analysis), and identifying the clusters of electrodes with a Monte Carlo *p*-value of less than .01. A more conservative alpha was used to account for additional variability introduced by combining the two patient subgroups. We then calculated the mean SW density, frequency, amplitude, upward slope, and downward slope of the electrodes within these clusters, which were used in subsequent analyses. In the first of these ROI analyses, we compared the frequency, amplitude, and slopes of the ROI electrodes across the two groups using linear mixed-effects models (SW Feature ∼ Group + Block + (1|Subject)). Then, we used multiple linear regressions to examine the relationship between each SW property that demonstrated significant between-group differences and two clinical variables: time since stroke and lesion volume. Note that these latter analyses were only conducted in a subset of *n*=21 stroke patients (*n*=11 left hemisphere, *n*=10 right hemisphere) for which we had imaging data that permitted estimation of lesion volume.

We then examined the effects of these stroke-related SWs on objective and subjective markers of behaviour within the stroke patients only. First, to examine the relationship between stroke-related SWs and perceptual decision-making, for each relevant SW property (as defined by the group-level difference analyses) we fitted LMMs: Hit Rate ∼ ROI SW Property + Block + (1|Subject) and RT ∼ ROI SW Property + Block + (1|Subject). Finally, we used linear regressions to examine whether these SW properties could predict either self-reported everyday functioning (score on the PCRS), or fatigue (FSS total score).

## Results

### Stroke is Associated with Slower Responses

Both groups of patients were slower to respond to contra- than ipsi-lesional stimuli (left stroke: 579.49 ± 41.15ms vs. 537.90 ± 36.02ms; right stroke: 782.84 ± 73.62ms vs. 650.65 ±26.66ms) in contrast with controls (left hemifield: 624.63 ± 28.68ms; right hemifield: 623.80 ± 23.58ms), as reported in Pearce et al (2025). Again, as reported previously, hit rate was broadly at ceiling across participants (left stroke: 93.63 ± 1.07%; right stroke: 90.52 ± 2.13%; control: 94.57 ± 0.32%). Importantly, when examining the relationship between speed (reaction times) and accuracy (hit rates) using LMEs, we found a negative correlation (Hit ∼ Block + RT + (1|Subject); t(488) = - 3.09, p = 0.002), which indicates that blocks in which participants slowed down led to higher accuracy. Examining the impact of the group revealed a significant interaction between RT and group (Hit ∼ Block*Group + RT + (1|Subject); F = 5.28, p = 0.02) and post-hoc comparisons showed a significant correlation between RT and Hit in the patient group (t(488) = -2.96, p = 0.003) but not in healthy controls (t(230) = 0.122, p = 0.9). These results evidence a speed-accuracy trade-off predominantly in patients, whereby faster responses were associated with poorer performance.

### Stroke is Associated with Increased Slow Wave (SW) Density

We then set out to compare SW density in the different groups. A model comparison (see Methods) showed no effect of the group (left hemisphere patients vs. right hemisphere patients vs. controls) on the threshold used to select SWs for each block (fixed effect of Group: F=0.02; p=0.88). We next examined the effect of block, electrode and group on SW Density (i.e., the number of SWs per minute) using a similar model comparison approach. The winning model comprised fixed effects of block, electrode and group as well as an interaction effect between electrode and block. Exploring this model, we found a significant effect of electrode (F=3.14, p<0.0001), group (F=5.60, p=0.018) and an interaction between electrode and block (F=1.69, p<0.001). The group effect was driven by a higher SW density for patients compared with controls (Fig. 1B). These results motivated the comparison of the scalp topographies of SW density across groups.

To examine topographical group differences in SW density, linear mixed-effects models (SW Density ∼ Group + Block + (1|Subject)) based on block-level estimates of SW density were fitted for each electrode. We also examined the difference within the patient group by splitting patients into left and right hemisphere stroke. We corrected for multiple comparisons using a cluster permutation approach with 500 permutations. Both stroke groups had significantly greater SW density across a majority of the scalp (Fig. 1B-C). We found no significant difference at any electrode between left and right stroke patients.

Together, these findings suggest that stroke patients demonstrate higher SW density than healthy controls across a majority of the scalp. Lesion hemisphere did not appear to meaningfully influence the distribution of SW detectable at the scalp.

### Region of Interest (ROI) Analyses

For all subsequent analyses within the stroke patients, we focused on a region of interest (ROI), that is a set of scalp electrodes over which SW properties were averaged. To achieve this, given the lack of significant differences between stroke groups we pooled all stroke patients and re-ran the above LMM with cluster permutations to compare control and patient SW densities across the scalp. We then defined a subset of electrodes (*N*=29) where SW density significantly differed from controls at the *α*=.01 level (as shown in Fig. 2A). Subsequent analyses were conducted by calculating mean values of interest across this subset of electrodes.

**Figure 2.**
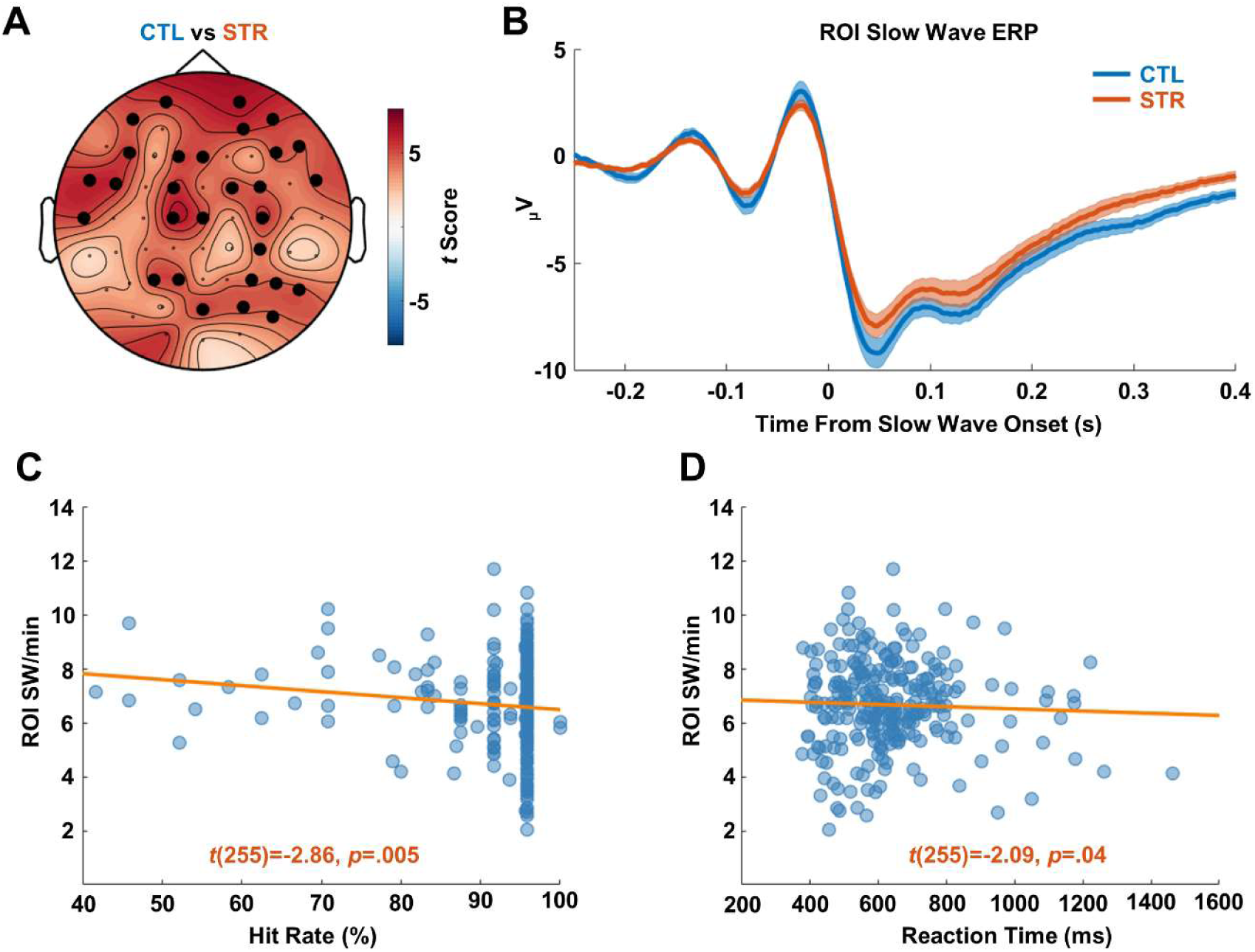
ROI Slow Wave (SW) Characteristics. *Note*. (A) Topographical maps of t-values denoting statistical slow wave (SW) density differences between controls and patients. Electrodes that form part of significant clusters (pcluster<.05) are denoted by bolded dots. CTL = control, STR = stroke. (B) Mean ERP within the defined ROI across all blocks, time-locked to SW onset, displayed separately for controls and patients. Also presented are scatter plots displaying the relationship between per-block ROI SW density and (C) hit rate and (D) RT. A least-squares regression line is fitted for reference.

### Regional Slow Waves (SWs) Share Similar Characteristics in Patients and Controls

We next sought to compare the spatiotemporal properties of the SWs within this ROI between controls and stroke patients. To this end, we first calculated block-level SW frequency (in Hz), peak-to-peak amplitude, and the gradient of the downward and upward slopes within the ROI. We then computed four LMMs with each of these variables as outcomes, with block and group as fixed effects and a random effect of subject (that is, equivalent models to those used for density). There was no significant difference between stroke patients and healthy controls on SW frequency (*t*(488)=0.69, *p*=.49), amplitude (*t*(488)=-0.80, *p*=.43), downward slope (*t*(488)=-1.51, *p*=.13), or upward slope (*t*(488)=-1.29, *p*=.20). Together, these results suggest that stroke does not alter the spatiotemporal properties of localised sleep-like SWs. Thus, we did not further analyse their effects on behaviour or functioning and instead focused on SW density. The average per-group SW waveform shown in Fig. 2B confirms this and shows that the temporal waveforms of SWs largely overlap across groups. Thus, SWs appear to differ across groups in terms of their rate of occurrence but not in terms of their properties.

### Higher Slow Wave (SW) Density is Associated with Poorer Task Performance

To examine the relationship between SWs and behaviour on the Bilateral Random Dot Motion (BRDM) task in stroke patients, we correlated the Hit Rate and RT per block with the SW density over the ROI (RT or Hit Rate ∼ ROI SW Density + Block + (1|Subject)). There was a significant negative relationship between ROI SW density and hit rate, *t*(255)=-2.86, *p*=.005, such that higher SW density predicted lower hit rate (Fig. 2C). There was also a significant relationship between higher ROI SW density and faster RT (*t*(255)=-2.09, *p*=.04; Fig. 2D). Importantly, since we showed a negative correlation between hit rate and RT in the patient group, a negative correlation between SW density and RT does not imply a gain in performance. Taken together, these results suggest that the emergence of post-stroke SWs is associated with modulations of perceptual reports during decision making.

### Relationship between Stroke-Related Slow Waves (SWs) and Everyday Functioning

We next aimed to identify whether SW density was associated with everyday functioning post-stroke. To this end we used two linear regressions to examine whether ROI SW density could predict either self-reported everyday functioning (PCRS), or fatigue (FSS).

We did not observe a significant relationship between SW density and everyday functioning scores (*t*(25)=-1.82, *p*=.08; Fig. 3A), although there was a non-significant trend in the expected direction (i.e., a negative correlation between SW density and everyday functioning), acknowledging the small sample size with which this analysis was conducted. There was no significant relationship between ROI SW density and fatigue scores (*t*(25)=0.16, *p*=.88; Fig. 3B).

**Figure 3.**
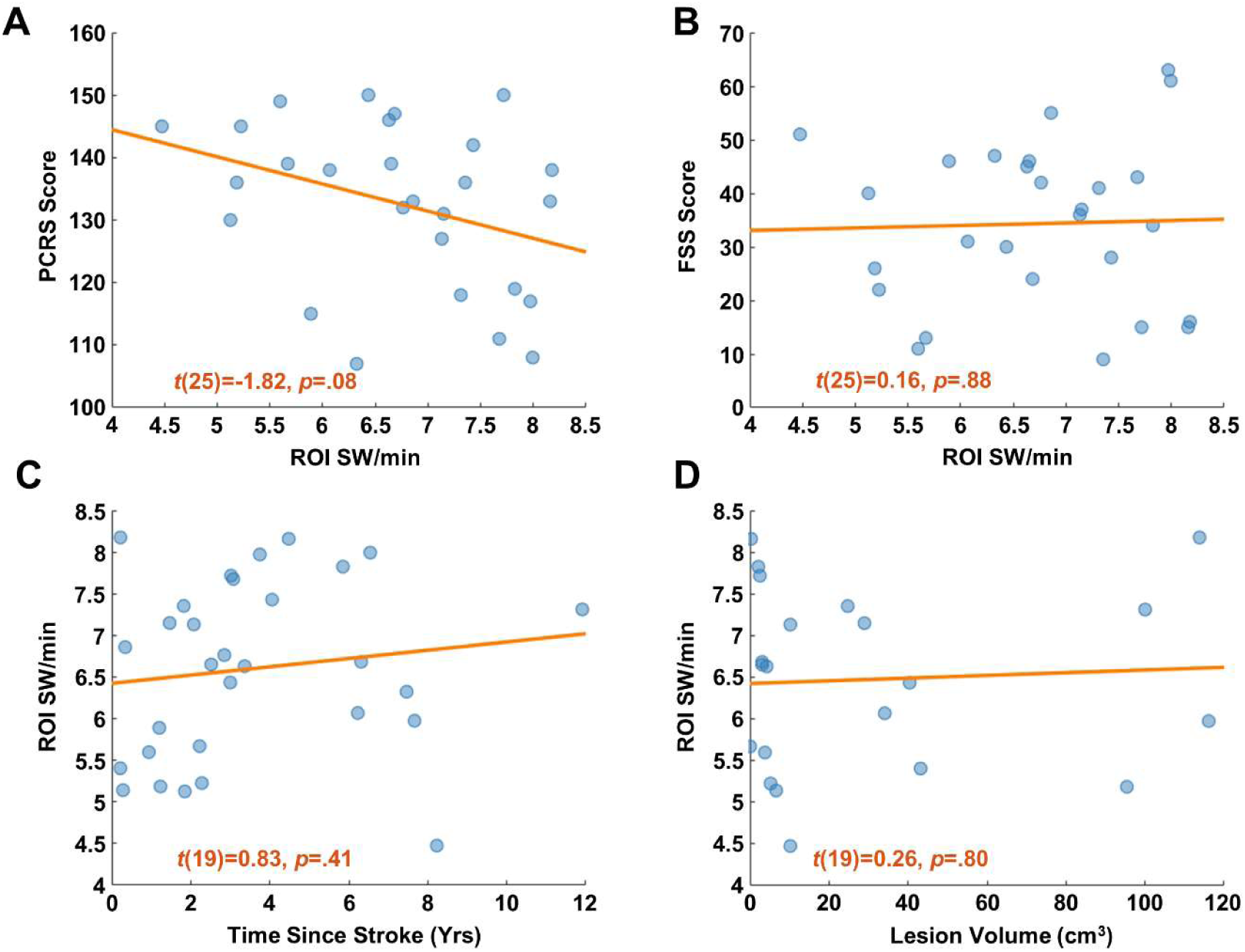
Regional Slow Wave (SW) Density Predicts Behavioural Performance. *Note.* Scatter plots depict the relationships between ROI slow wave (SW) density and (A) PCRS score, (B) FSS score, (C) time since stroke, and (D) lesion volume. A least-squares regression line is fitted for reference. T-values and p-values were computed with the LMMs described in the Methods.

### Stroke-Related Slow Wave (SW) Density Effects are not an Artefact of Lesion Characteristics

Next, in our stroke patients only we sought to understand whether stroke characteristics (i.e., lesion volume and time since stroke) were predictive of ROI SW density using two linear multiple regressions, with age included as a covariate. We did not find evidence of a relationship between ROI SW density and either time since stroke, *t*(19)=0.83, *p*=.41 (Fig. 3C), or lesion volume, *t*(19)=0.26, *p*=.80 (Fig. 3D).

## Discussion

Here, we used EEG and a perceptual decision-making task to examine the occurrence of sleep-like SWs in chronic stroke patients and their relationship with behavioural performance. Sleep-like SWs shared similar spatiotemporal properties across groups but occurred more frequently in stroke patients than in healthy controls. This effect was prominent scalp-wide, with a similar distribution regardless of the hemisphere of the lesion. Additionally, we showed evidence linking SWs with an alteration of the accuracy and speed of decisions in a perceptual task. To our knowledge, this represents the first empirical evidence suggesting a relationship between sleep-like SWs and post-stroke cognitive impairments.

Stroke-related increases in sleep-like SW density were observed at most scalp sites and were similarly distributed regardless of lesion hemisphere, size, or recency of stroke, suggesting a generalised increase in SW activity across the scalp. The increase in SW density was also observed across blocks. These results complement previous findings reporting an increase in such SWs following external perturbations (TMS or electrical stimulation (Pigorini et al., 2015; Sarasso et al., 2020)), demonstrating that passive and non-invasive recordings can also reveal this shift toward sleep-like dynamics. Yet, the lack of spatial specificity appears in contrast to TMS-EEG findings showing that SWs are maximally generated following activation of the perilesional area, and solely in the ipsi-rather than contra-lesional hemisphere (Sarasso et al., 2020). This discrepancy could be related to the difference between the observation of spontaneous waves (as in our case) and the induction of SWs through external perturbations. Moreover, while appearing to arise from perilesional tissue, post-lesion SWs have been observed in regions remote from the lesion, even via intracranial recordings (Buchkremer-Ratzmann et al., 1996; Rorden & Karnath, 2004; Russo et al., 2021). This occurrence of SWs far from the lesioned area could reflect a phenomenon of diaschisis (Feeney & Baron, 1986; Massimini et al., 2024), whereby brain regions connected to the lesioned area also show changes in cortical dynamics. Thus, in addition to standard volume conduction and the lack of spatial resolution of scalp EEG, our findings could also reflect the propagation of SWs from perilesional tissue across structurally intact brain networks via cortico-cortical and/or subcortico-cortical pathways (Massimini et al., 2024; Russo et al., 2021). Finally, another possibility is that the spatially generalised pattern that emerged here is attributable to the fact that we tested chronic, rather than acute stroke patients, such that non-lesioned but interconnected neurons may develop an increased propensity to fall into off-periods over time. This may be expected due to the recruitment of other networks to compensate for dysfunctional tissue (Frost et al., 2003; Wang et al., 2010). These findings thus point to the need for longitudinal efforts to track the spatiotemporal evolution of SWs over the acute-to-chronic period, with a view to clarifying the cause, purpose, and mechanisms of sleep-like SW propagation following stroke.

Our results build upon past work by establishing a previously unexplored relationship between post-lesion sleep-like SWs and poor performance on perceptual tasks. Previous studies showed a positive link between the occurrence of SWs after a stroke and recovery (Finnigan et al., 2004, 2007; Sarasso et al., 2024), which might be linked to the homeostatic functions of these SWs during sleep. Indeed, sleep SWs are indeed involved in the maintenance of metabolic or synaptic homeostasis (Tononi & Cirelli, 2014; Wisor et al., 2013), a role that could be partially preserved when occurring during wakefulness (Driessen et al., 2025; Krueger et al., 2019; Yang et al., 2025). In rodents, the induction of OFF periods via optogenetics in post-stroke asleep mice promoted plasticity and improved functional recovery (Facchin et al., 2020). Yet, this role of post-stroke SWs in assisting with future recovery could be at the cost of cognitive functioning (Massimini et al., 2024). Indeed, our findings suggest that these restorative benefits come at a potential cost of in-the-moment functioning. A relationship between the intrusion of sleep-like SWs and attentional errors has previously been demonstrated in healthy adults (Andrillon et al., 2021; Bernardi et al., 2015; Hung et al., 2013; Pinggal et al., 2022; Quercia et al., 2018), and is consistent with the well-known negative impact of declining arousal on selective attention and task performance (Robbins, 1997; Unsworth & Robison, 2018). Furthermore, it has been suggested that sleep-like SWs may index the core mechanism by which undamaged regions remote to the lesion show dysfunction (diaschisis) (Feeney & Baron, 1986), such that underlying neuronal off-periods result in significant network-level interference that has widespread consequences for cognition (Massimini et al., 2024; Russo et al., 2021). The link between sleep-like SWs and broader dysfunction thus presents an intriguing avenue for future research.

The mechanistic relationship between the occurrence of SWs and behavioural impairments could also be indirect. Another plausible explanation is that this relationship may be mediated by the degree of patient recovery as opposed to representing a direct causal perturbation of cognitive processes associated with decision-making. Post-lesion sleep-like SWs decrease in accordance with the normalisation of functional networks and clinical recovery (Sarasso et al., 2024). Those with higher SW density may therefore have a higher need for ongoing restorative processes, and thus be more likely to show behavioural impairment. Clarification could be achieved in studies with higher trial counts that enable behavioural performance comparisons between trials with and without SW intrusions, thus providing clearer insight into whether the occurrence of SWs contemporaneously disrupts behavioural function. An intriguing further question to be answered by larger studies is whether variation in lesion location alters the behavioural consequences of sleep-like SWs, as may be suggested by the moderating influence of spatial location on the relationship between SWs and behaviour in healthy adults (Andrillon et al., 2021). For example, investigations of whether SWs emerging from one hemisphere are more detrimental for contra- rather than ipsi-lateral perceptual responding would enable greater appreciation of the network-wide effects of SW intrusion. Examining the link between lesions, SW occurrence and behaviour in larger populations and more varied cognitive tasks would allow to explore this question.

A final possibility is that our findings suggest heightened daytime sleepiness in our stroke patients. Given that stroke is associated with high rates of sleep disturbance (Hermann & Bassetti, 2009; Iddagoda et al., 2020) and heightened sleep pressure increases the intrusion of SWs in wakefulness (Hung et al., 2013; Rector et al., 2009; Vyazovskiy et al., 2011), increased SWs in our stroke cohort could also reflect a poorer sleep quality as opposed to direct lesion effects. This raises the further possibility that poorer task performance with increasing SW density could be mediated by sleep quality, via the known relationship between poor sleep and attentional dysregulation (Lim & Dinges, 2010). Yet, we did not find evidence that sleep-like SWs were associated with self-reported post-stroke fatigue, although sleepiness and fatigue are independent constructs (Shen et al., 2006). Future studies that seek to disentangle the sleep-like SWs that arise from stroke-related sleep disturbance from those that directly result from damaged tissue may provide a clearer picture of whether they intrude upon functioning. Direct interventions on sleep could help not only to examine the relationship between SWs and cognitive deficits but also to test new therapeutic strategies to mitigate these symptoms (e.g., (Weightman et al., 2024)).

In summary, here we leveraged an integrated EEG/perceptual decision-making paradigm to investigate the density and behavioural effects of sleep-like SWs following stroke. We showed that passive, non-invasive EEG recordings evidence an increase in sleep-like SWs following stroke. In addition, we demonstrate for the first time that increased intrusion of sleep-like SWs in stroke patients is associated with altered reporting of perceptual stimuli. Collectively, we add weight to the growing body of evidence for sleep-like SWs as a reliable measure of localised brain arousal states and their effect on cognition. Pending future large-scale longitudinal work to better characterise the direct or indirect relationship between sleep-like SWs and behaviour, sleep-like SWs present as a plausible mechanism of post-lesion functional network dysfunction and subsequent cognitive deficits.

## References

1. Andrillon, T., Burns, A., MacKay, T., Windt, J., & Tsuchiya, N. (2021). Predicting lapses of attention with sleep-like slow waves. Nature Communications. 10.1101/2020.06.23.166991

2. Andrillon, T., & Kouider, S. (2019). The vigilant sleeper: Neural mechanisms of sensory (de)coupling during sleep. Current Opinion in Physiology. 10.1016/j.cophys.2019.12.002

3. Andrillon, T., & Oudiette, D. (2023). What is sleep exactly? Global and local modulations of sleep oscillations all around the clock. Neuroscience & Biobehavioral Reviews, 155, 105465. 10.1016/j.neubiorev.2023.105465

4. Andrillon, T., Windt, J., Silk, T., Drummond, S. P. A., Bellgrove, M. A., & Tsuchiya, N. (2019). Does the Mind Wander When the Brain Takes a Break? Local Sleep in Wakefulness, Attentional Lapses and Mind-Wandering. Frontiers in Neuroscience, 13. 10.3389/fnins.2019.00949

5. Avvenuti, G., Bertelloni, D., Lettieri, G., Ricciardi, E., Cecchetti, L., Pietrini, P., & Bernardi, G. (2021). Emotion Regulation Failures Are Preceded by Local Increases in Sleep-like Activity. Journal of Cognitive Neuroscience, 33(11), 2342–2356. 10.1162/jocn_a_01753

6. Barker-Collo, S., Feigin, V. L., Parag, V., Lawes, C. M. M., & Senior, H. (2010). Auckland Stroke Outcomes Study: Part 2: Cognition and functional outcomes 5 years poststroke. Neurology, 75(18), 1608–1616. 10.1212/WNL.0b013e3181fb44c8

7. Barskova, T., & Wilz, G. (2006). Psychosocial functioning after stroke: Psychometric properties of the patient competency rating scale. Brain Injury, 20(13–14), 1431–1437. 10.1080/02699050600976317

8. Bernardi, G., Siclari, F., Yu, X., Zennig, C., Bellesi, M., Ricciardi, E., Cirelli, C., Ghilardi, M. F., Pietrini, P., & Tononi, G. (2015). Neural and Behavioral Correlates of Extended Training during Sleep Deprivation in Humans: Evidence for Local, Task-Specific Effects. Journal of Neuroscience, 35(11), 4487–4500. 10.1523/JNEUROSCI.4567-14.2015

9. Brainard, D. H. (1997). The Psychophysics Toolbox. Spatial Vision, 10(4), 433–436.

10. Britten, K. H., Shadlen, M. N., Newsome, W. T., & Movshon, J. A. (1992). The analysis of visual motion: A comparison of neuronal and psychophysical performance. J.Neurosci., 12(12), 4745–4765.

11. Buchkremer-Ratzmann, I., August, M., Hagemann, G., & Witte, O. W. (1996). Electrophysiological Transcortical Diaschisis After Cortical Photothrombosis in Rat Brain. Stroke, 27(6), 1105–1111. 10.1161/01.STR.27.6.1105

12. Calderon, D. P., Kilinc, M., Maritan, A., Banavar, J. R., & Pfaff, D. (2016). Generalized CNS arousal: An elementary force within the vertebrate nervous system. Neuroscience & Biobehavioral Reviews, 68, 167–176. 10.1016/j.neubiorev.2016.05.014

13. Carson, N., Leach, L., & Murphy, K. J. (2018). A re-examination of Montreal Cognitive Assessment (MoCA) cutoff scores. International Journal of Geriatric Psychiatry, 33(2), 379–388. 10.1002/gps.4756

14. Chaudhuri, A., & Behan, P. O. (2004). Fatigue in neurological disorders. The Lancet, 363(9413), 978–988. 10.1016/S0140-6736(04)15794-2

15. Chen, W., Jiang, T., Huang, H., & Zeng, J. (2023). Post-stroke fatigue: A review of development, prevalence, predisposing factors, measurements, and treatments. Frontiers in Neurology, 14, 1298915. 10.3389/fneur.2023.1298915

16. Colombo, M. A., Favaro, J., Mikulan, E., Pigorini, A., Zauli, F. M., Sartori, I., d’Orio, P., Castana, L., Toldo, I., Sartori, S., Sarasso, S., Bayne, T., Seth, A. K., & Massimini, M. (2025). Hemispherotomy leads to persistent sleep-like slow waves in the isolated cortex of awake humans. PLOS Biology, 23(10), e3003060. 10.1371/journal.pbio.3003060

17. Cornelissen, F. W., Peters, E. M., & Palmer, J. (2002). The Eyelink Toolbox: Eye tracking with MATLAB and the Psychophysics Toolbox. Behavior Research Methods, Instruments, & Computers, 34(4), 613–617. 10.3758/BF03195489

18. Driessen, K., Squarcio, F., Tononi, G., & Cirelli, C. (2025). Induction of cortical ON/OFF periods in awake mice fulfills sleep functions. 10.1101/2025.10.04.680459

19. Eason, R. G., Harter, M. R., & White, C. T. (1969). Effects of attention and arousal on visually evoked cortical potentials and reaction time in man. Physiology & Behavior, 4(3), 283–289. 10.1016/0031-9384(69)90176-0

20. Facchin, L., Schöne, C., Mensen, A., Bandarabadi, M., Pilotto, F., Saxena, S., Libourel, P. A., Bassetti, C. L. A., & Adamantidis, A. R. (2020). Slow Waves Promote Sleep-Dependent Plasticity and Functional Recovery after Stroke. The Journal of Neuroscience: The Official Journal of the Society for Neuroscience, 40(45), 8637–8651. 10.1523/JNEUROSCI.0373-20.2020

21. Feeney, D. M., & Baron, J. C. (1986). Diaschisis. Stroke, 17(5), 817–830. 10.1161/01.STR.17.5.817

22. Finnigan, S. P., Rose, S. E., Walsh, M., Griffin, M., Janke, A. L., McMahon, K. L., Gillies, R., Strudwick, M. W., Pettigrew, C. M., Semple, J., Brown, J., Brown, P., & Chalk, J. B. (2004). Correlation of Quantitative EEG in Acute Ischemic Stroke With 30-Day NIHSS Score: Comparison With Diffusion and Perfusion MRI. Stroke, 35(4), 899–903. 10.1161/01.STR.0000122622.73916.d2

23. Finnigan, S. P., Walsh, M., Rose, S. E., & Chalk, J. B. (2007). Quantitative EEG indices of sub-acute ischaemic stroke correlate with clinical outcomes. Clinical Neurophysiology, 118(11), 2525–2532. 10.1016/j.clinph.2007.07.021

24. Frost, S. B., Barbay, S., Friel, K. M., Plautz, E. J., & Nudo, R. J. (2003). Reorganization of Remote Cortical Regions After Ischemic Brain Injury: A Potential Substrate for Stroke Recovery. Journal of Neurophysiology, 89(6), 3205–3214. 10.1152/jn.01143.2002

25. Hermann, D. M., & Bassetti, C. L. (2009). Sleep-related breathing and sleep-wake disturbances in ischemic stroke. Neurology, 73(16), 1313–1322. 10.1212/WNL.0b013e3181bd137c

26. Hung, C.-S., Sarasso, S., Ferrarelli, F., Riedner, B., Ghilardi, M. F., Cirelli, C., & Tononi, G. (2013). Local experience-dependent changes in the wake EEG after prolonged wakefulness. Sleep, 36(1), 59–72. 10.5665/sleep.2302

27. Hyvärinen, A., & Oja, E. (2000). Independent component analysis: Algorithms and applications. Neural Networks, 13(4–5), 411–430. 10.1016/S0893-6080(00)00026-5

28. Iddagoda, M. T., Inderjeeth, C. A., Chan, K., & Raymond, W. D. (2020). Post-stroke sleep disturbances and rehabilitation outcomes: A prospective cohort study. Internal Medicine Journal, 50(2), 208–213. 10.1111/imj.14372

29. Jokinen, H., Melkas, S., Ylikoski, R., Pohjasvaara, T., Kaste, M., Erkinjuntti, T., & Hietanen, M. (2015). Post-stroke cognitive impairment is common even after successful clinical recovery. European Journal of Neurology, 22(9), 1288–1294. 10.1111/ene.12743

30. Krueger, J. M., Nguyen, J. T., Dykstra-Aiello, C. J., & Taishi, P. (2019). Local sleep. Sleep Medicine Reviews, 43, 14–21. 10.1016/j.smrv.2018.10.001

31. Krupp, L. B. (1989). The Fatigue Severity Scale: Application to Patients With Multiple Sclerosis and Systemic Lupus Erythematosus. Archives of Neurology, 46(10), 1121. 10.1001/archneur.1989.00520460115022

32. Kusec, A., Milosevich, E., Williams, O. A., Chiu, E. G., Watson, P., Carrick, C., Drozdowska, B. A., Dillon, A., Jennings, T., Anderson, B., Dawes, H., Thomas, S., Kuppuswamy, A., Pendlebury, S. T., Quinn, T. J., & Demeyere, N. (2023). Long-term psychological outcomes following stroke: The OX-CHRONIC study. BMC Neurology, 23(1), 426. 10.1186/s12883-023-03463-5

33. Lazar, R. M., Fitzsimmons, B.-F., Marshall, R. S., Berman, M. F., Bustillo, M. A., Young, W. L., Mohr, J. P., Shah, J., & Robinson, J. V. (2002). Reemergence of Stroke Deficits With Midazolam Challenge. Stroke, 33(1), 283–285. 10.1161/hs0102.101222

34. Léger, D., Debellemaniere, E., Rabat, A., Bayon, V., Benchenane, K., & Chennaoui, M. (2018). Slow-Wave Sleep: From the Cell to the Clinic. Sleep Medicine Reviews. 10.1016/j.smrv.2018.01.008

35. Lim, J., & Dinges, D. F. (2010). A meta-analysis of the impact of short-term sleep deprivation on cognitive variables. Psychological Bulletin, 136(3), 375–389. 10.1037/a0018883

36. Losier, B. J. W., & Klein, R. M. (2001). A review of the evidence for a disengage deficit following parietal lobe damage. Neuroscience & Biobehavioral Reviews, 25(1), 1–13. 10.1016/S0149-7634(00)00046-4

37. Loughnane, G. M., Newman, D. P., Bellgrove, M. A., Lalor, E. C., Kelly, S. P., & O’Connell, R. G. (2016). Target Selection Signals Influence Perceptual Decisions by Modulating the Onset and Rate of Evidence Accumulation. Current Biology, 26(4), 496–502. 10.1016/j.cub.2015.12.049

38. Maris, E., & Oostenveld, R. (2007). Nonparametric statistical testing of EEG- and MEG-data. Journal of Neuroscience Methods, 164(1), 177–190. 10.1016/j.jneumeth.2007.03.024

39. Marmelshtein, A., Eckerling, A., Hadad, B., Ben-Eliyahu, S., & Nir, Y. (2023). Sleep-like changes in neural processing emerge during sleep deprivation in early auditory cortex. Current Biology, 33(14), 2925–2940.e6. 10.1016/j.cub.2023.06.022

40. Massimini, M., Corbetta, M., Sanchez-Vives, M. V., Andrillon, T., Deco, G., Rosanova, M., & Sarasso, S. (2024). Sleep-like cortical dynamics during wakefulness and their network effects following brain injury. Nature Communications, 15(1), 7207. 10.1038/s41467-024-51586-1

41. Massimini, M., Ferrarelli, F., Esser, S. K., Riedner, B. A., Huber, R., Murphy, M., Peterson, M. J., & Tononi, G. (2007). Triggering sleep slow waves by transcranial magnetic stimulation. Proc Natl Acad Sci U S A, 104, 8496–8501. 10.1073/pnas.0702495104

42. Murphy, M., Huber, R., Esser, S., A. Riedner, B., Massimini, M., Ferrarelli, F., Felice Ghilardi, M., & Tononi, G. (2011). The Cortical Topography of Local Sleep. Current Topics in Medicinal Chemistry, 11(19), 2438–2446. 10.2174/156802611797470303

43. Murri, L., Gori, S., Massetani, R., Bonanni, E., Marcella, F., & Milani, S. (1998). Evaluation of acute ischemic stroke using quantitative EEG: A comparison with conventional EEG and CT scan. Neurophysiologie Clinique/Clinical Neurophysiology, 28(3), 249–257. 10.1016/S0987-7053(98)80115-9

44. Nadarajah, M., Mazlan, M., Abdul-Latif, L., & Goh, H. (2017). Test-retest reliability, internal consistency and concurrent validity of Fatigue Severity Scale in measuring post-stroke fatigue. European Journal of Physical and Rehabilitation Medicine, 53(5). 10.23736/S1973-9087.16.04388-4

45. Nasreddine, Z. S., Phillips, N. A., Bédirian, V., Charbonneau, S., Whitehead, V., Collin, I., Cummings, J. L., & Chertkow, H. (2005). The Montreal Cognitive Assessment, MoCA: A Brief Screening Tool For Mild Cognitive Impairment. Journal of the American Geriatrics Society, 53(4), 695–699. 10.1111/j.1532-5415.2005.53221.x

46. Newsome, W. T., Britten, K. H., & Movshon, J. A. (1989). Neuronal correlates of a perceptual decision. Nature, 341, 52–54.

47. Nuwer, M. R. (1996). Quantitative EEG analysis in clinical settings. Brain Topography, 8(3), 201–208. 10.1007/BF01184770

48. Oostenveld, R., Fries, P., Maris, E., & Schoffelen, J.-M. (2011). FieldTrip: Open source software for advanced analysis of MEG, EEG, and invasive electrophysiological data. Computational Intelligence and Neuroscience, 2011, 156869. 10.1155/2011/156869

49. Ozyemisci-Taskiran, O., Batur, E. B., Yuksel, S., Cengiz, M., & Karatas, G. K. (2019). Validity and reliability of fatigue severity scale in stroke. Topics in Stroke Rehabilitation, 26(2), 122–127. 10.1080/10749357.2018.1550957

50. Pearce, D. J., Loughnane, G. M., Chong, T. T.-J., Demeyere, N., Mattingley, J. B., Moore, M. J., New, P. W., O’Connell, R. G., O’Neill, M. H., Rangelov, D., Stolwyk, R. J., Webb, S. S., Zhou, S.-H., Brosnan, M. B., & Bellgrove, M. A. (2023). Neurophysiological mechanisms underlying post-stroke deficits in contralesional perceptual processing. 10.1101/2023.12.12.571233

51. Pelli, D. G. (1997). The VideoToolbox software for visual psychophysics: Transforming numbers into movies. Spatial Vision, 10(4), 437–442.

52. Pendlebury, S. T., & Rothwell, P. M. (2019). Incidence and prevalence of dementia associated with transient ischaemic attack and stroke: Analysis of the population-based Oxford Vascular Study. The Lancet Neurology, 18(3), 248–258. 10.1016/S1474-4422(18)30442-3

53. Petersen, A., Petersen, A. H., Bundesen, C., Vangkilde, S., & Habekost, T. (2017). The effect of phasic auditory alerting on visual perception. Cognition, 165, 73–81. 10.1016/j.cognition.2017.04.004

54. Pfaff, D., & Banavar, J. R. (2007). A theoretical framework for CNS arousal. BioEssays, 29(8), 803–810. 10.1002/bies.20611

55. Pfaff, D., Ribeiro, A., Matthews, J., & Kow, L. (2008). Concepts and Mechanisms of Generalized Central Nervous System Arousal. Annals of the New York Academy of Sciences, 1129(1), 11–25. 10.1196/annals.1417.019

56. Pfaff, D. W., & Kieffer, B. L. (2008). Preface. Annals of the New York Academy of Sciences, 1129(1). 10.1196/annals.1417.034

57. Pigorini, A., Sarasso, S., Proserpio, P., Szymanski, C., Arnulfo, G., Casarotto, S., Fecchio, M., Rosanova, M., Mariotti, M., Lo Russo, G., Palva, J. M., Nobili, L., & Massimini, M. (2015). Bistability breaks-off deterministic responses to intracortical stimulation during non-REM sleep. NeuroImage, 112, 105–113. 10.1016/j.neuroimage.2015.02.056

58. Pinggal, E., Dockree, P. M., O’Connell, R. G., Bellgrove, M. A., & Andrillon, T. (2022). Pharmacological manipulations of physiological arousal and sleep-like slow waves modulate sustained attention. The Journal of Neuroscience, JN-RM-0836–22. 10.1523/JNEUROSCI.0836-22.2022

59. Price, C. J., Hope, T. M., & Seghier, M. L. (2017). Ten problems and solutions when predicting individual outcome from lesion site after stroke. NeuroImage, 145, 200–208. 10.1016/j.neuroimage.2016.08.006

60. Prigatano, G. P., & Neuropsychological Rehabilitation Program (Presbyterian Hospital, O. C., Okla.). (1986). Neuropsychological rehabilitation after brain injury. Johns Hopkins University Press. https://cir.nii.ac.jp/crid/1971712334726418976

61. Purdon, P. L., Pierce, E. T., Mukamel, E. A., Prerau, M. J., Walsh, J. L., Wong, K. F. K., Salazar-Gomez, A. F., Harrell, P. G., Sampson, A. L., Cimenser, A., Ching, S., Kopell, N. J., Tavares-Stoeckel, C., Habeeb, K., Merhar, R., & Brown, E. N. (2013). Electroencephalogram signatures of loss and recovery of consciousness from propofol. Proceedings of the National Academy of Sciences, 110(12). 10.1073/pnas.1221180110

62. Quercia, A., Zappasodi, F., Committeri, G., & Ferrara, M. (2018). Local Use-Dependent Sleep in Wakefulness Links Performance Errors to Learning. Frontiers in Human Neuroscience, 12. 10.3389/fnhum.2018.00122

63. Rector, D. M., Schei, J. L., Van Dongen, H. P. A., Belenky, G., & Krueger, J. M. (2009). Physiological markers of local sleep. European Journal of Neuroscience, 29(9), 1771–1778. 10.1111/j.1460-9568.2009.06717.x

64. Rector, D. M., Topchiy, I. A., Carter, K. M., & Rojas, M. J. (2005). Local functional state differences between rat cortical columns. Brain Research, 1047(1), 45–55. 10.1016/j.brainres.2005.04.002

65. Riedner, B. A., Vyazovskiy, V. V., Huber, R., Massimini, M., Esser, S., Murphy, M., & Tononi, G. (2007). Sleep homeostasis and cortical synchronization: III. A high-density EEG study of sleep slow waves in humans. Sleep, 30, 1643–1657.

66. Robbins, T. W. (1997). Arousal systems and attentional processes. Biological Psychology, 45(1–3), 57–71. 10.1016/S0301-0511(96)05222-2

67. Robertson, I. H., Mattingley, J. B., Rorden, C., & Driver, J. (1998). Phasic alerting of neglect patients overcomes their spatial deficit in visual awareness. Nature, 395(6698), 169–172. 10.1038/25993

68. Rogasch, N. C., Thomson, R. H., Farzan, F., Fitzgibbon, B. M., Bailey, N. W., Hernandez-Pavon, J. C., Daskalakis, Z. J., & Fitzgerald, P. B. (2014). Removing artefacts from TMS-EEG recordings using independent component analysis: Importance for assessing prefrontal and motor cortex network properties. NeuroImage, 101, 425–439. 10.1016/j.neuroimage.2014.07.037

69. Rorden, C., & Karnath, H.-O. (2004). Using human brain lesions to infer function: A relic from a past era in the fMRI age? Nature Reviews Neuroscience, 5(10), 812–819. 10.1038/nrn1521

70. Rosanova, M., Fecchio, M., Casarotto, S., Sarasso, S., Casali, A. G., Pigorini, A., Comanducci, A., Seregni, F., Devalle, G., Citerio, G., Bodart, O., Boly, M., Gosseries, O., Laureys, S., & Massimini, M. (2018). Sleep-like cortical OFF-periods disrupt causality and complexity in the brain of unresponsive wakefulness syndrome patients. Nature Communications, 9(1), 4427. 10.1038/s41467-018-06871-1

71. Russo, S., Pigorini, A., Mikulan, E., Sarasso, S., Rubino, A., Zauli, F. M., Parmigiani, S., d’Orio, P., Cattani, A., Francione, S., Tassi, L., Bassetti, C. L. A., Lo Russo, G., Nobili, L., Sartori, I., & Massimini, M. (2021). Focal lesions induce large-scale percolation of sleep-like intracerebral activity in awake humans. NeuroImage, 234, 117964. 10.1016/j.neuroimage.2021.117964

72. Sarasso, S., D’Ambrosio, S., Fecchio, M., Casarotto, S., Viganò, A., Landi, C., Mattavelli, G., Gosseries, O., Quarenghi, M., Laureys, S., Devalle, G., Rosanova, M., & Massimini, M. (2020). Local sleep-like cortical reactivity in the awake brain after focal injury. Brain, 143(12), 3672–3684. 10.1093/brain/awaa338

73. Sarasso, S., D’Ambrosio, S., Russo, S., Bernardelli, L., Hassan, G., Comanducci, A., De Giampaulis, P., Dalla Vecchia, L., Lanzone, J., & Massimini, M. (2024). The reduction of sleep-like perilesional cortical dynamics underlies clinical recovery in stroke. 10.1101/2024.03.16.24304272

74. Sexton, E., McLoughlin, A., Williams, D. J., Merriman, N. A., Donnelly, N., Rohde, D., Hickey, A., Wren, M.-A., & Bennett, K. (2019). Systematic review and meta-analysis of the prevalence of cognitive impairment no dementia in the first year post-stroke. European Stroke Journal, 4(2), 160–171. 10.1177/2396987318825484

75. Shen, J., Barbera, J., & Shapiro, C. M. (2006). Distinguishing sleepiness and fatigue: Focus on definition and measurement. Sleep Medicine Reviews, 10(1), 63–76. 10.1016/j.smrv.2005.05.004

76. Sheybani, L., Vivekananda, U., Rodionov, R., Diehl, B., Chowdhury, F. A., McEvoy, A. W., Miserocchi, A., Bisby, J. A., Bush, D., Burgess, N., & Walker, M. C. (2023). Wake slow waves in focal human epilepsy impact network activity and cognition. Nature Communications, 14(1), 7397. 10.1038/s41467-023-42971-3

77. Steriade, M., Dossi, R., & Nunez, A. (1991). Network modulation of a slow intrinsic oscillation of cat thalamocortical neurons implicated in sleep delta waves: Cortically induced synchronization and brainstem cholinergic suppression. The Journal of Neuroscience, 11(10), 3200–3217. 10.1523/JNEUROSCI.11-10-03200.1991

78. Stolwyk, R. J., Mihaljcic, T., Wong, D. K., Chapman, J. E., & Rogers, J. M. (2021). Poststroke Cognitive Impairment Negatively Impacts Activity and Participation Outcomes: A Systematic Review and Meta-Analysis. Stroke, 52(2), 748–760. 10.1161/STROKEAHA.120.032215

79. Stolwyk, R. J., Mihaljcic, T., Wong, D. K., Hernandez, D. R., Wolff, B., & Rogers, J. M. (2024). Post-stroke Cognition is Associated with Stroke Survivor Quality of Life and Caregiver Outcomes: A Systematic Review and Meta-analysis. Neuropsychology Review, 34(4), 1235–1264. 10.1007/s11065-024-09635-5

80. Tononi, G., & Cirelli, C. (2014). Sleep and the Price of Plasticity: From Synaptic and Cellular Homeostasis to Memory Consolidation and Integration. Neuron, 81(1), 12–34. 10.1016/j.neuron.2013.12.025

81. Tononi, G., & Massimini, M. (2008). Why does consciousness fade in early sleep? Annals of the New York Academy of Sciences, 1129, 330–334. 10.1196/annals.1417.024

82. Unsworth, N., & Robison, M. K. (2018). Tracking arousal state and mind wandering with pupillometry. *Cognitive, Affective*, & Behavioral Neuroscience, 18(4), 638–664. 10.3758/s13415-018-0594-4

83. Vyazovskiy, V. V., & Harris, K. D. (2013). Sleep and the single neuron: The role of global slow oscillations in individual cell rest. Nature Reviews Neuroscience, 14(6), 443–451. 10.1038/nrn3494

84. Vyazovskiy, V. V., Olcese, U., Hanlon, E. C., Nir, Y., Cirelli, C., & Tononi, G. (2011). Local sleep in awake rats. Nature, 472, 443–447. 10.1038/nature10009

85. Vyazovskiy, V. V., Olcese, U., Lazimy, Y. M., Faraguna, U., Esser, S. K., Williams, J. C., Cirelli, C., & Tononi, G. (2009). Cortical firing and sleep homeostasis. Neuron, 63, 865–878.

86. Wang, L., Yu, C., Chen, H., Qin, W., He, Y., Fan, F., Zhang, Y., Wang, M., Li, K., Zang, Y., Woodward, T. S., & Zhu, C. (2010). Dynamic functional reorganization of the motor execution network after stroke. Brain, 133(4), 1224–1238. 10.1093/brain/awq043

87. Webb, S. S., Hobden, G., Roberts, R., Chiu, E. G., King, S., & Demeyere, N. (2022). Validation of the UK English Oxford cognitive screen-plus in sub-acute and chronic stroke survivors. European Stroke Journal, 7(4), 476–486. 10.1177/23969873221119940

88. Weightman, M., Robinson, B., Mitchell, M. P., Garratt, E., Teal, R., Rudgewick-Brown, A., Demeyere, N., Fleming, M. K., & Johansen-Berg, H. (2024). Sleep and motor learning in stroke (SMiLES): A longitudinal study investigating sleep-dependent consolidation of motor sequence learning in the context of recovery after stroke. BMJ Open, 14(2), e077442. 10.1136/bmjopen-2023-077442

89. Wisor, J. P., Rempe, M. J., Schmidt, M. A., Moore, M. E., & Clegern, W. C. (2013). Sleep Slow-Wave Activity Regulates Cerebral Glycolytic Metabolism. Cerebral Cortex, 23(8), 1978–1987. 10.1093/cercor/bhs189

90. Yang, Z., Williams, S. D., Beldzik, E., Anakwe, S., Schimmelpfennig, E., & Lewis, L. D. (2025). Attentional failures after sleep deprivation are locked to joint neurovascular, pupil and cerebrospinal fluid flow dynamics. Nature Neuroscience. 10.1038/s41593-025-02098-8

91. Yokoyama, E., Nagata, K., Hirata, Y., Satoh, Y., Watahiki, Y., & Yuya, H. (1996). Correlation of EEG activities between slow-wave sleep and wakefulness in patients with supra-tentorial stroke. Brain Topography, 8(3), 269–273. 10.1007/BF01184783

